# PyRice: a Python package for querying *Oryza Sativa* databases

**DOI:** 10.1101/2020.04.20.049742

**Authors:** Quan Do, Ho Bich Hai, Pierre Larmande

## Abstract

**Summary:** Currently, gene information available for *Oryza sativa* species is located in various online heterogeneous data sources. Moreover, methods of access are also diverse, mostly web-based and sometimes query APIs, which might not always be straightforward for domain experts. The challenge is to collect information quickly from these applications and combine it logically, to facilitate scientific research. We developed a Python package named PyRice, a unified programming API to access all supported databases at the same time with consistent output. PyRice design is modular and implements a smart query system which fits the computing resources to optimize the query speed. As a result, PyRice is easy to use and produces intuitive results.

**Availability and implementation:** https://github.com/SouthGreenPlatform/PyRice

**Documentation:** https://pyrice.readthedocs.io

**Licence information:** MIT

**Supplementary information:** Supplementary data are available online.

## 1 Introduction

Rice, a model crop plant, is a major cereal grain widely consumed by a large part of the world’s human population, especially in Asia. Since two decades, many digital resources have been developed in rice genomics. However, compared to human genomics, not as much centralized resources and analysis tools are available for rice genomics. Moreover, most of the information currently available are scattered and patchy in nature. Thus, for scientists, the challenge lies in integrating data and finding useful information. In the scope of the project, we aim to build an API to solve the problem of collecting and managing gene and gene products information from different sources. The PyRice package is developed to run remote queries over ten databases and web applications so far. However, PyRice provides generic wrappers to extend this number with more datatypes such as genetics, gene expression and pathways, according (Garg P. *et al*., 2016) databases classification. Moreover, PyRice uses parallel processing to improve query speed. Furthermore, it indexes results for a fast search and also supports exporting results into different formats.

## 2 Materials and Methods

Information of *Oryza sativa* genes are published on several open-access databases using different gene annotation models, e.g. RAPDB (Hiroaki *et al*., 2013), MSU7 or dedicated IDs (i.e. SNP-SEEK (Mansueto *et al*., 2016) and IC4R (IC4R Project Consortium *et al*., 2016). PyRice manages a dictionary of ID mapping across databases since each uses either of the two systems RAPDB and MSU7 (e.g. LOC_Os01g01010 = Os01g0100100; while the first ID is from MSU7 and the second is from RAPDB). Supplementary Fig. S1 shows the flowchart of the package. There are two main query types: with genomic coordinates or with a list of gene IDs.

### 2.1 Databases

Currently, PyRice supports 10 databases to query gene information: Oryzabase (Kurata *et al*., 2006), RAPDB, Funricegenes (Yao *et al*., 2017), Gramene (Tello-Ruiz *et al*., 2018), IC4R, MSU7 Rice GAP, Rice SNP-SEEK, EMBL-EBI Expression Atlas (Papatheodorou *et al*., 2017) and GWAS Atlas (Tian *et al*., 2019).

In PyRice, databases information is stored in a unique XML configuration file which allows the users to easily add new databases or update information. In order to help with connection failure due to network or database server’s unavailability, some lightweight databases (i.e. SNP-SEEK, Oryzabase, RAPDB, GWAS Atlas) are downloaded and cached locally in PyRice after their first query. They are regularly updated within package by built-in function. The PyRice package builds a cross-reference dictionary between three types of index (i.e. MSU7, RAPDB and SNP-SEEK) to facilitate information retrieval. Essentially, this means PyRice creates a dictionary of ID mappings with the ones mentioned above, valid them and assign them with internal IDs. It also contains for each gene, ID mappings along with genomic coordinates extracted from SNP-SEEK database.

### 2.2 Parallel processing

We implemented a smart query system which fits the computing resources and manages the query processing (presented in section 2.3). PyRice runs independently each task in parallel, thus it is able to ignore interrupted tasks due to internet error correction or databases maintenance which may lead to conflict between queries. Moreover, we implemented several internet client authentication methods in order to avoid server rejections. Since computing resources could have different architectures, PyRice implements 3 parallel options (i.e. multi-processing, multi-threading and the combination of two). Thus, users can choose the number of threads and CPUs depending on their computer resource. In Supplementary File 1, we evaluated the benefits of using the parallel architectures.

### 2.3 Query system

In PyRice, the query system is based on a database wrapper model. Because online databases usually provide web UI or APIs to query information, we developed a dozen of wrappers for each type of querying methods (e.g. GET, POST), input parameters, and output formats. Furthermore, it is possible to configure some filters to keep the necessary attributes. Results are indexed by IDs and stored in dictionaries along with databases information (presented in section 2.4). As mentioned previously, *Oryza sativa* databases use diverse types of IDs to store information. PyRice handles it with a cross-reference dictionary of ID mappings. There are two types of query that is supported in PyRice, based on this dictionary:

- Using IDs: PyRice automatically detects the type of query IDs and maps to those used in the requested database. ID availability check is also conducted before executing the query.
- Using genomic coordinates: These are defined by chromosome number, start and end position.

### 2.4 Results

As presented in section 2.2, PyRice stores results in a dictionary structure (shown in Supplementary File 2). Since gene information exists in several databases, this allows organizing information quickly while saving computing resources. This dictionary is then used to produce final results. Currently, the supported outputs are:

- A summary file is created for each gene found in the initial query. This file gathers all information extracted from the databases and uses the HTML or JSON format for convenient browsing (see Supplementary Fig. S3);
- A spreadsheet stores the resulting gene list along with basic information. PyRice supports output formats of CSV, JSON, HTML, and pickle (see Supplementary Fig. S2).

Pickle format (i.e. .pkl) is a serializing Python object structure which scales up very well with distributed processes, hence can handle a large number of gene queries. PyRice also provides a search and filtering functions for this format, i.e. complex searches using SQL logical operators, i.e. AND, OR, NOT, and result filtering.

## 3. Conclusion

PyRice is a python package that simplifies querying *Oryza sativa* databases, making it easier to synthesize gene information. PyRice allows multiple processes that help to increase the speed of queries when multi-core processors are available. The usage is straightforward, producing intuitive results. It can be easily extended with new databases (see Supplementary File 3), potentially applying to other plants as well.

In the near future, we will extend PyRice to support more databases and new species. Besides, we will provide user support to add new users’ databases. To speed up the query process, which is currently parallelized on a single computer, a distributed version of PyRice will be developed. Thus, the new version will use the resources of HPC clusters.

## Acknowledgements

The authors thank the former students, Dang Lam and Vautrin Baptiste who implemented the first prototype of this package.

