## Supplementary File 3 for "PyRice: a Python package for querying *Oryza Sativa* databases"

Also available at https://pyrice.readthedocs.io/en/latest/pyrice_instruction.html#add-new-database

### Add new database

PyRice package supports queries on new databases by adding its description manually in *database_description.xml*. Using JSON format, here is an example with SNP-SEEK database API: https://snp-seek.irri.org/ws/genomics/gene/osnippo/chr01?start=1&end=15000&model=iric:

```
<database dbname="snpseek" type="text/JSON" method="GET" normalize="false">
    <link stern="https://snp-seek.irri.org/ws/genomics/gene/osnippo/" aft=""/>
    <fields>
        <field></field>
        <field op="=">start</field>
        <field op="=">end</field>
        <field op="=">model</field>
    </fields>
</database>
```

#### For more details:

- dbname : database name
- type : the result returned by API
- method : GET/POST (default GET)
- normalize : normalize name of database true/false (default false)
- stern : URL of API
- op : parameters (see on API above)

For example, with an API from Planteome: http://browser.planteome.org/api/search/annotation?bioentity=AT4G32150:

```
<database dbname="planteome" type="text/JSON" method="GET" normalize="false">
    <link stern="http://browser.planteome.org/api/search/annotation?"
aft=""></link>
    <fields>
        <field op="=">bioentity</field>
    </fields>
</database>
```
