## Supplementary File 2 for "PyRice: a Python package for querying *Oryza Sativa* databases"

```
Output database:
{'OsNippo01g010050': {
    'rapdb': {
        'Locus_ID': 'Os01g0100100',
        'Description': 'RabGAP/TBC domain containing protein.',
            'Oryzabase Gene Name Synonym(s)': 'Molecular Function: Rab GTPase
activator activity (GO:0005097)',
            ...},
        'gramene': {
            '_id': 'Os01g0100100',
            'name': 'Os01g0100100',
            'biotype': 'protein_coding',
            ...},
        ...},
    'OsNippo01g010150': {
        'rapdb': {...},
        'gramene': {...},
        ...},
    ...
}
```

```
Output database:
{'OsNippo01g010050': {
        'rapdb': {
            'Locus_ID': 'Os01g0100100',
            'Description': 'RabGAP/TBC domain containing protein.',
            'Position': '',
            ...},
        'ic4r': {
            'Anther_Normal': {'expression_value': '0.699962'},
            'Anther_WT': {'expression_value': '13.9268'},
            ...},
        ...},
    'OsNippo01g010300': {
        'rapdb': {...},
        'ic4r': {...},
        ...},
    ...
}
```
Examples of dictionary structures used to store results in PyRice.
