## Supplementary File 1 for "PyRice: a Python package for querying *Oryza Sativa* databases"

We performed test queries with a set of genes on several databases and presented the results in the online documentation, along with a tutorial. In addition, we evaluated the query performance of PyRice as shown in table below. The benchmark was implemented on a server with Intel(R) Xeon(R) CPU E5-2683 v3 @ 2.00GHz having 56 cores, 112 threads and 128GB RAM. We ran several tests with different gene set sizes and various combinations of processing types. As the table shows, on a set of 10,000 genes, it took 530 seconds with multi-processing and multi-threading options.

Generally, it is feasible to use PyRice for querying a large number of genes with a standard laptop (i.e. ~4 cores). However, the Internet connectivity and the web server response that could affect the performance, it is, thus, recommended to use multiple CPU cores, if available.

**Table 1. Query performance evaluation of PyRice**

| Case | Number process | 500 genes | 1000 genes | 10000 genes |
|---|---|---|---|---|
| Case 1 | | 1175s | 2569s | 22085s |
| Case 2 | 4 | 306s | 656s | 5741s |
| | 8 | 168s | 342s | 2861s |
| | 16 | 100s | 186s | 1510s |
| Case 3 | 4 | 360s | 792s | 6583s |
| | 8 | 273s | 598s | 5418s |
| | 16 | 280s | 612s | 5488s |
| Case 4 | 4 | 105s | 210s | 1803s |
| | 8 | 60s | 115s | 955s |
| | 16 | 43s | 70s | 530s |

Case 1: Do not use parallel processing;
Case 2: Using only multi-processing (Number of process is the number of core running, one process opens one thread);
Case 3: Using only multi-threading (Number of process is the number of threads running, it uses only one core);
***Case 4: Using both multi-processing and multi-threading (Number of process is the number of core running, by default one process opens two threads).***
