## Supplementary Fig 3 for "PyRice: a Python package for querying *Oryza Sativa* databases"

| Database | Field | Value |
|---|---|---|
| oryzabase | **Trait Gene Id** | 22606 |
| | **CGSNL Gene Symbol** | TLP27 |
| | **Gene symbol synonym(s)** | OsTLP27 |
| | **CGSNL Gene Name** | THYLAKOID LUMENAL PROTEIN 27 |
| | **Gene name synonym(s)** | Thylakoid lumenal protein 27 |
| | **Protein Name** | THYLAKOID LUMENAL PROTEIN 27 |
| | **Explanation** | AtTLP homolog. TO:0020116: photochemical quenching. |
| | **Trait Class** | Coloration - Chlorophyll |
| | **RAP ID** | Os01g0102300 |
| | **MUS ID** | LOC_Os01g01280.1 |
| | **Gene Ontology** | GO:0015979 - photosynthesis, GO:0009658 - chloroplast organization, GO:0043476 - pigment accumulation, GO:0007623 - circadian rhythm, GO:0009543 - chloroplast thylakoid lumen |
| | **Trait Ontology** | TO:0000494 - pigment content, TO:0002715 - chloroplast development trait |
| | **Plant Ontology** | PO:0025034 - leaf , PO:0020104 - leaf sheath |
| rapdb | **Transcript_ID** | Os01t0102300-01 |
| | **Locus_ID** | Os01g0102300 |
| | **Description** | Thylakoid lumen protein, Photosynthesis and chloroplast development |
| | **RAP-DB Gene Symbol Synonym(s)** | OsTLP27 |
| | **CGSNL Gene Symbol** | TLP27 |
| | **CGSNL Gene Name** | THYLAKOID LUMENAL PROTEIN 27 |
| | **Oryzabase Gene Symbol Synonym(s)** | OsTLP27 |
| | **Oryzabase Gene Name Synonym(s)** | Thylakoid lumenal protein 27 |
| | **Transcript evidence** | AK067320 |
| | **ORF evidence** | Q94CX1 |
| | **Curation Date** | March 7, 2017 |
| | **Literature PMID** | 22921006 |

*Figure 3. The gene information results output in HTML format. The filename correspond to the SNP-SEEK gene ID. Each database have a row section containing information retrieved from the query.*
