## Supplementary Fig 2 for "PyRice: a Python package for querying *Oryza Sativa* databases"

### Supplementary Fig. 1

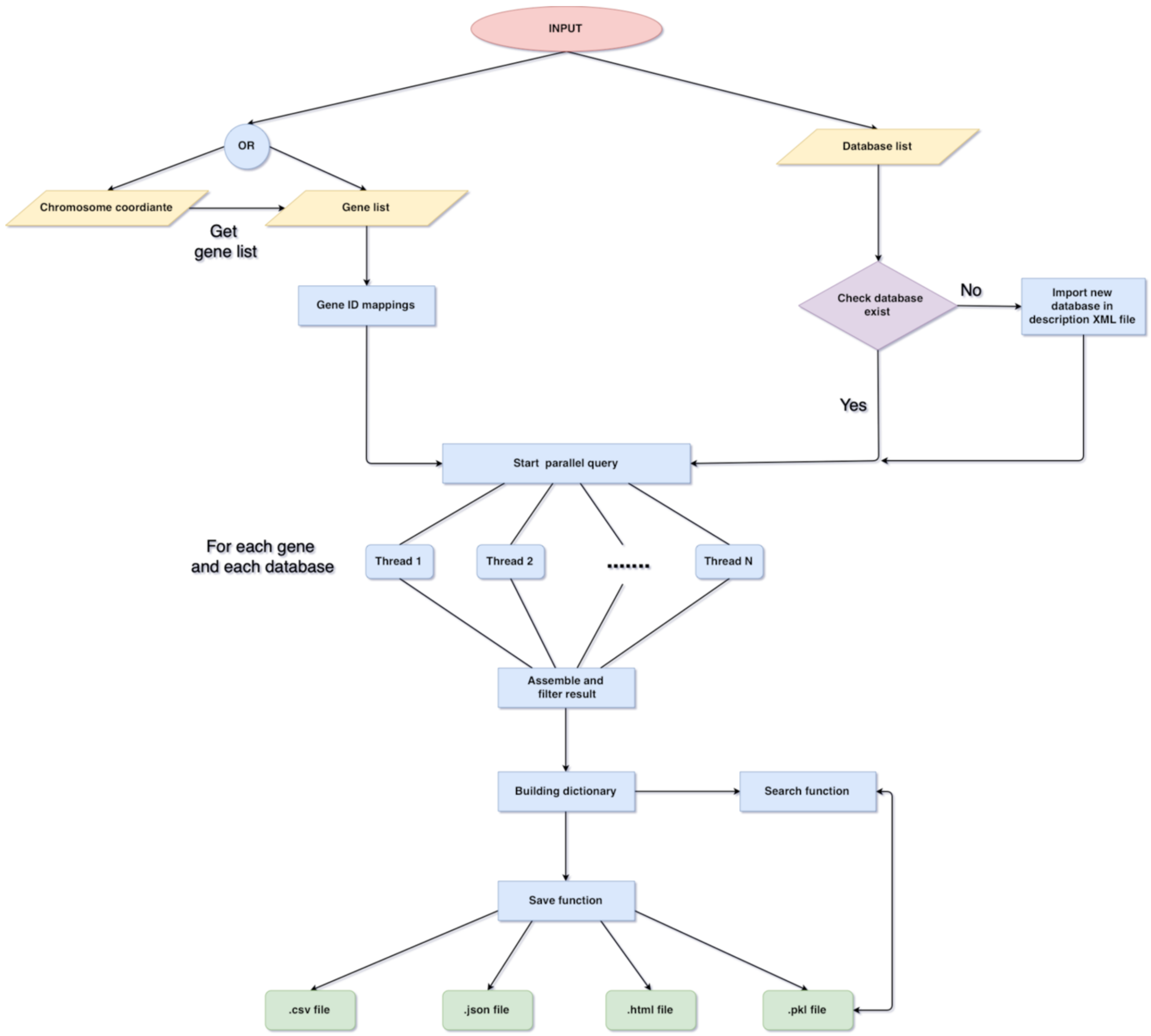


*Figure 1. The flowchart of the PyRice starts with the input of genomic coordinates or gene identifiers, and a list of databases. PyRice automatically processes the queries and saves the results in the requested formats.*
