## Supplementary File 1 for "PyRice: a Python package for querying *Oryza Sativa* databases"

### Supplementary Fig. 2

| | rapdb.Locus_ID | rapdb.Description | rapdb.Oryzabase Gene Name Synonym(s) | gramene._id | gramene.name | gramene.description | gramene.biotype | gramene.taxon_id | gramene.system_name | gramene.db_type | gramene.gene_idx | gramene.location |
|---|---|---|---|---|---|---|---|---|---|---|---|---|
| OsNippo01g010050 | Os01g0100100 | RabGAP/TBC domain containing protein. | Molecular Function: Rab GTPase activator activ... | Os01g0100100 | Os01g0100100 | RabGAP/TBC domain containing protein. (Os01t01... | protein_coding | 39947.0 | oryza_sativa | core | 0.0 | {'region': '1', 'start': 2983, 'end': 10815, '... |
| OsNippo01g010100 | Os01g0100300 | Cytochrome P450 domain containing protein. | Biological Process: oxidation-reduction proces... | Os01g0100300 | Os01g0100300 | Cytochrome P450 domain containing protein. (Os... | protein_coding | 39947.0 | oryza_sativa | core | 2.0 | {'region': '1', 'start': 11372, 'end': 12284, ... |
| OsNippo01g010150 | Os01g0100200 | Conserved hypothetical protein. | NaN | Os01g0100200 | Os01g0100200 | Conserved hypothetical protein. (Os01t0100200-01) | protein_coding | 39947.0 | oryza_sativa | core | 1.0 | {'region': '1', 'start': 11218, 'end': 12435, ... |
| OsNippo01g010200 | Os01g0100400 | Similar to Pectinesterase-like protein. | Biological Process: oxidation-reduction proces... | Os01g0100400 | Os01g0100400 | Similar to Pectinesterase-like protein. (Os01t... | protein_coding | 39947.0 | oryza_sativa | core | 3.0 | {'region': '1', 'start': 12721, 'end': 15685, ... |

*Figure 2 : Example of table results output in HTML format. Each gene information is summarized in one row. Gene IDs are linked to their detailed output ( see Supplementary Fig. 3) and listed in the first column.*
